# A self-renewing biomimetic skeletal muscle construct engineered using induced myogenic progenitor cells

**DOI:** 10.1101/2023.04.10.534929

**Authors:** Inseon Kim, Seunghun S. Lee, Adhideb Ghosh, Stephen J. Ferguson, Ori Bar-Nur

## Abstract

Skeletal muscle is a highly organized and regenerative tissue that maintains its homeostasis primarily by activation and differentiation of muscle stem cells. Mimicking an *in vitro* skeletal muscle differentiation program that contains self-renewing adult muscle stem cells and aligned myotubes has been challenging. Here, we set out to engineer a biomimetic skeletal muscle construct that can self-regenerate and produce aligned myotubes using induced myogenic progenitor cells (iMPCs), a heterogeneous culture consisting of skeletal muscle stem, progenitor and differentiated cells. Utilizing electrospinning, we fabricated polycaprolactone (PCL) substrates that enabled iMPC-differentiation into aligned myotubes by controlling PCL fiber orientation. Newly-conceived constructs contained highly organized multinucleated myotubes in conjunction with self-renewing muscle stem cells, whose differentiation capacity was augmented by Matrigel supplementation. Additionally, we demonstrate using single cell RNA-sequencing (scRNA-seq) that iMPC-derived constructs faithfully recapitulate a step-wise myogenic differentiation program. Notably, when the constructs were subjected to a damaging myonecrotic agent, self-renewing muscle stem cells rapidly differentiated into aligned myotubes, akin to skeletal muscle repair *in vivo*. Taken together, we report on a novel *in vitro* system that mirrors myogenic regeneration and muscle fiber alignment, and can serve as a platform to study myogenesis, model muscular dystrophies or perform drug screens.

## Introduction

Body movements are governed by voluntary contractions of multinucleated muscle fibers (myofibers), the functional component of skeletal muscle tissue^1^. Due to its role in generating locomotion, skeletal muscle tissue displays a unique architecture comprised of highly organized myofibers, which enable muscles to contract and produce movement in a desired direction^1^. In response to injury or disease state, skeletal muscle tissue can rapidly regenerate by resident stem cells termed satellite cells (SCs)^2,3^. These cells reside between the basal lamina layer and the sarcolemma, and are typically found in a quiescent state primed for activation^2^. Furthermore, SCs express high level of the transcription factor paired box 7 (PAX7) and exhibit elevated activity of the Notch signaling pathway^4,5^. Upon insult, SCs play a pivotal role in restoring homeostasis by forming new myofibers or repairing damaged ones^2,6^. During this process, quiescent SCs (QSCs) are activated to form a proliferative cell population that can either self-renew and return to quiescence, or differentiate into committed progenitors and fusion-competent myocytes that merge with damaged myofibers for tissue repair^2,6^.

The isolation of SCs from muscles and their serial passaging *in vitro* as proliferative myoblasts has been central to the study of myogenesis^7,8^. Typically, primary or immortalized myoblasts such as C2C12 are cultured in a high serum condition and differentiate into myotubes by serum withdrawal, thus permitting the study of skeletal muscle formation *in vitro*^7,8^. However, this assay typically gives rise to disorganized myotubes in the culture dish, rendering tissue engineering a highly desirable approach to produce aligned and functional skeletal muscle constructs^9-11^. Indeed, in past decades a multitude of studies reported on ordered architecture construction of skeletal muscle cells by using a variety of fabrication techniques including 3D printing^12^, micropatterning^13^, acoustic patterning^14,15^, magnetic actuation^16^, filamented light-beam biofabrication^17^ and electrospinning^11,18-20^. These efforts were harnessed to produce organized muscle constructs, however most often reported only on production of aligned myotubes in the absence of self-renewing muscle stem cells, thus limiting the study of myogenesis in biomimetic constructs^11,20^.

Electrospinning is a high-throughput scaffold fabrication technique that enables production of nanofiber membrane apt for cell growth^21^. This technique facilitates orientation control of electrospun nano-scaled fibers on the substrate, guiding cellular growth akin to 3D bioprinting^18^. As a consequence of these attributes, electrospun substrates have been utilized to produce highly aligned myotubes using multiple materials including decellularized extracellular matrix (ECM)^22,23^, synthetic or natural polymers such as polyurethane^24^, poly(ε-caprolactone) (PCL)^25^, poly(lactic-co-glycolic acid)^26^, fibrin/alginate^27^ and gelatin^11,28^. However, incorporation of self-renewing muscle stem cells into electrospun nano-scaled fibers that can regenerate muscle fibers has rarely been reported^10,11^.

To date, a variety of myogenic cells have been used for skeletal muscle tissue engineering purposes, predominantly immortalized myoblasts such as C2C12 or primary myoblasts^9-11^. Yet, these myogenic cells do not faithfully mirror proliferating and self-renewing SCs and myoblasts tend to lose differentiation capacities with extended passaging^29^. As an alternative, several studies have used pluripotent stem cell (PSC)-derived myogenic precursors, which share cellular features with muscle stem cells and can differentiate into myotubes^30-34^. However, PSC-derived myogenic precursors and derivative myotubes may carry embryonic attributes, and require intricate differentiation protocols which can give rise to heterogeneous cultures containing multiple cell types^33,35,36^.

An alternative myogenic cell source suitable for skeletal muscle engineering encompasses induced myogenic progenitor cells (iMPCs), which are directly reprogramed from fibroblasts by myogenic transcription factors, oftentimes in the presence of small molecules^37-40^. Several recent studies reported on production of iMPCs by short-lived expression of the canonical myogenic transcription factor MyoD in conjunction with the small molecules Forskolin, RepSox and CHIR99021 (F/R/C)^37,41,42^. Notably, iMPC cultures are composed of disorganized multinucleated and contractile myotubes, in addition to self-renewing muscle stem cells that can expand extensively while maintaining differentiation capacity^37^. Furthermore, unlike conventional myoblasts, muscle stem cells present in iMPCs share several attributes with *in vivo* proliferating SCs, thus serving as a potential superior cell source for skeletal muscle engineering^41,42^. Moreover, iMPCs require a relatively short time to produce and demonstrate cellular heterogeneity of regenerating skeletal muscle cells^41^. However, whether iMPCs produced with MyoD and F/R/C supplementation can recapitulate a myogenic differentiation program in a 3D microenvironment and produce aligned myotubes, similar to muscle fibers *in vivo*, has not been investigated. Herein we set out to engineer an aligned skeletal muscle construct by growing iMPCs on electrospun PCL substrates, and further investigated whether the incorporation of Matrigel (MA), a potent and widely used biomaterial, can augment the differentiation capacity of iMPCs. Additionally, we utilized scRNA-seq of the constructs to dissect cell populations and lineage trajectories indicative of a myogenic regeneration program. Last, we subjected the constructs to serial cardiotoxin injury assay and assessed the potential of iMPCs to regrow aligned myotubes following insult, akin to regeneration *in vivo*.

## Results

### Engineering and characterizing iMPC-derived skeletal muscle constructs (iSMCs)

We set out to generate biomimetic skeletal muscle constructs consisting of aligned myotubes and muscle stem cells in fabricated scaffolds, generated via electrospinning of PCL nanofibers (Figure 1). By controlling the speed of the rotating collector during electrospinning, two types of electrospun PCL scaffolds were obtained: an aligned pattern (AP) at high speed (1500rpm) and a random pattern (RP) at low speed (100rpm) (Figure 1). As the next step, we seeded onto the scaffolds previously generated iMPCs in the presence of iMPC medium containing F/R/C to promote cell expansion and differentiation, purposing to engineer aligned iSMCs resembling skeletal muscle (Figure 1)^41^.

**Figure 1.**
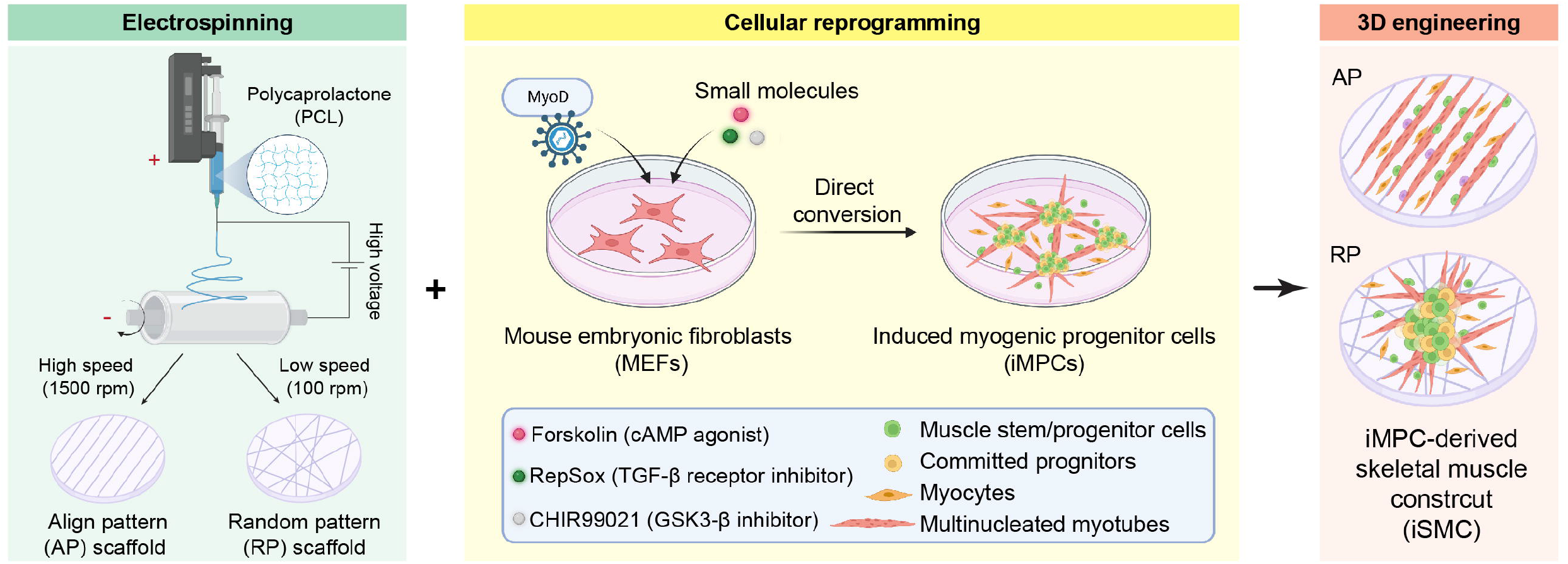
A schematic illustration of experimental design. The schematic illustrates the goal of this study: engineering skeletal muscle constructs by seeding fibroblast-derived induced myogenic progenitor cells (iMPCs) onto electrospun PCL scaffolds containing either aligned pattern (AP) or random pattern (RP) nanofibers.

We first confirmed by Field Emission-Scanning Electron Microscopy (FE-SEM) the unidirectional pattern of the nanofibers in AP scaffolds, whereas a disorganized and non-aligned pattern was formed in RP scaffolds (Figure 2A). To quantify the alignment degree of the nanofibers, we measured their angle orientation and observed significant differences between AP and RP scaffolds (58.2±18.2° and 12.8±5.3° from the point of origin, respectively) (Figures 2A and 2B). Of note, the diameter of electrospun nanofibers in the AP scaffold was smaller due to increased tension elicited by high-speed electrospinning (Figures 2A and S1A). Next, to facilitate a non-toxic environment prior to cell seeding, the scaffolds were treated with Sodium Hydroxide (NaOH) for 4 hours (hrs), followed by overnight incubation with 70% ethanol and sterilization by UV light for 30 minutes (mins). The NaOH treatment was used to increase scaffold hydrophilicity by hydrating ester bonds in the PCL polymers, thereby exposing the carboxyl groups on the surface^43^. In addition, MA was incorporated into a few scaffolds to increase the biocompatibility and assess the growth of iMPCs when cultured in the presence of potent ECM proteins. This combined effort revealed that under all 4 conditions (AP, RP, AP+MA and RP+MA), a biocompatible environment for iMPC growth was established as observed by FE-SEM (Figure 2C). Strikingly, iMPCs seeded onto AP and AP+MA scaffolds were highly aligned, growing in a unidirectional pattern dictated by the orientation of the PCL nanofibers, while iMPCs cultured on RP and RP+MA scaffolds exhibited a disorganized orientation pattern (Figure 2C).

**Figure 2.**
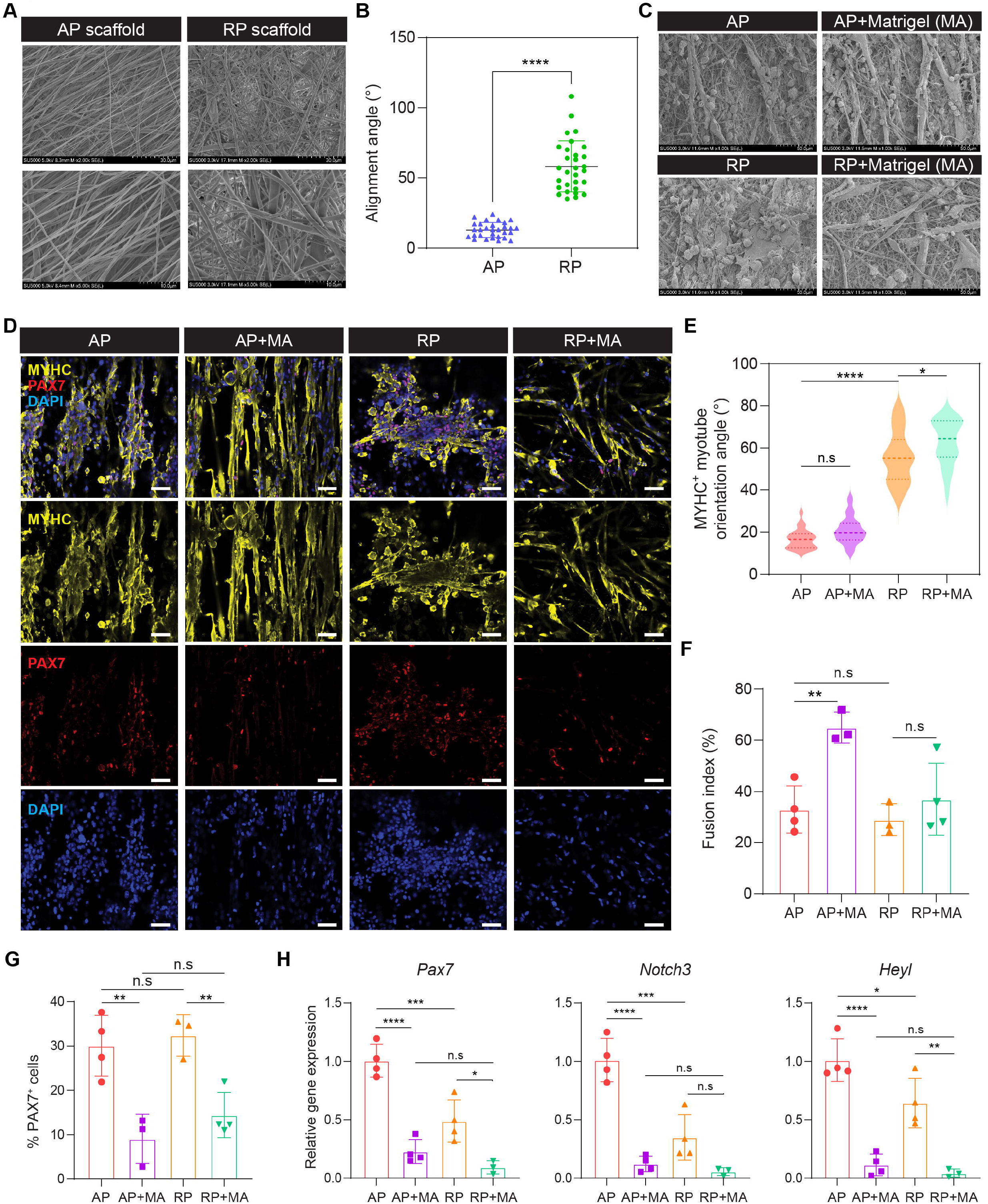
Molecular characterization of iSMCs. **(A)** Electron microscopy images of electrospun PCL scaffolds showing either an aligned pattern (AP) or random pattern (RP). Images were acquired by Field Emission-scanning Electron Microscopy (FE-SEM). **(B)** Angle measurement of PCL electrospun fibers shown in (A). The data are shown as means□±□S.D (n=30). Statistical significance was determined by a two-tailed unpaired t-test (****p<0.0001). **(C)** FE-SEM images of iMPCs seeded onto electrospun PCL scaffolds in the indicated conditions at day 2 after cell seeding. **(D)** Representative immunofluorescence images of iSMCs stained for MYHC (yellow), PAX7 (red), and DAPI (blue) for the respective conditions. Scale bar, 50 µm. **(E)** A violin plot showing the alignment angle degree of representative MYHC positive myotubes as shown in (D) (n=30). Statistical significance was determined by one-way ANOVA (*p<0.5, ****p<0.0001, n.s = nonsignificant). **(F)** Graph showing fusion index of myotubes, calculated as the percentage of myonuclei in multinucleated myotubes with more than 2 nuclei over all nuclei in an examined field. The data are own as means□±□S.D (n=3-4 independent experiments). Statistical significance was determined by a one-way ANOVA (**p<0.01, n.s = nonsignificant). **(G)** Graph showing the quantification of the percentage of PAX7 positive cells over the total number of DAPI positive cells in iSCMs under the conditions demonstrated in (D). The data are shown as means□±□S.D (n=4 independent experiments). Statistical significance was determined by a one-way ANOVA (**p<0.01, n.s = nonsignificant). **(H)** Quantitative RT-PCR of the indicated myogenic stem cells markers. The data are shown as means□±□S.D (n=3-4 independent experiments). Statistical significance was determined by a one-way ANOVA (*p<0.5, **p<0.01, ***p<0.001, ****p<0.0001, n.s = nonsignificant).

To molecularly characterize iSMCs, we immunostained the constructs for the differentiation marker myosin heavy chain (MYHC) and the muscle stem cell marker PAX7, documenting positive cells for either protein under all conditions (Figures 2D and S1B). In agreement with the FE-SEM analysis, MYHC^+^ myotubes detected under AP or AP+MA conditions were well-aligned, exhibiting smaller alignment angle in comparison with either the RP or RP+MA conditions (Figures 2D, 2E and S1B). Furthermore, MYHC^+^ myotubes generated under AP+MA displayed a significantly higher fusion index in respect to AP, suggesting elevated differentiation capacity of iMPCs into myotubes in the presence of MA (Figure 2F). Furthermore, under AP and RP conditions, a higher number of PAX7^+^ cells were recorded than under the respective conditions containing MA (around 30% vs. 10%) (Figure 2G). This result was further confirmed by elevated gene expression of other SC markers (*Notch3* and *Heyl*) under AP and RP conditions in comparison to the respective MA-supplemented groups (Figure 2H). However, the expression of several commitment myogenic genes was not significantly different between the various conditions, aside from *Myh1* under the RP+MA condition (Figure S1C). Taken together, aligned iSMCs were generated by growing iMPCs in biofabricated PCL scaffolds. These constructs consisted of PAX7^+^ stem cells as well as aligned MYHC^+^ myotubes, whose differentiation capacity was augmented by MA supplementation.

### Dissecting proliferation dynamics and myogenic cell populations in iSMCs

Skeletal muscle differentiation initiates by non-proliferative SCs that undergo activation into proliferative progenitors termed myoblasts and fusion-competent myocytes^2^. Given the presence of both multinucleated MYHC^+^ myotubes and PAX7^+^ stem cells in iSMCs, we next opted to characterize the proliferation dynamics and differentiation capacities of myogenic cell populations in iSMCs (Figure 3A). To this end, we performed a 5-Ethynyl-2’-deoxyuridine (EdU) analysis in concert with immunostaining for MYOD, a myoblast and differentiation cell marker (Figure 3B). In accordance with the PAX7 immunostaining, this analysis revealed a higher percentage of EdU^+^ cells under AP or RP conditions (20.7±5.7% and 17.9±4.8%, respectively) in comparison to the AP+MA and RP+MA conditions (9.7±3.9% and 6.9±2.3%, respectively) (Figures 3B and 3C). This observation further supports the notion that MA supplementation elicits cell cycle arrest and differentiation into post-mitotic cells. Of note, among the 4 conditions there were no major differences in the number of MYOD^+^ cells, as in all iSMCs between 80-90% MYOD^+^ cells were detected (Figures 3B and 3D). Further investigation revealed that in the 4 conditions, around 70% of the cells were MYOD^+^/EdU^-^ and as such non-cycling cells, whereas between 7-20% were MYOD^+^/EdU^+^ cycling cells (Figures 3E and S2A). Interestingly, under AP and RP conditions, a higher percentage of MYOD^+^/EdU^+^ proliferative cells and lower MYOD^+^/EdU^-^ non-proliferative cells were detected in comparison to their respective MA-treated conditions, suggesting increased proliferation (Figures 3E and S2A).

**Figure 3.**
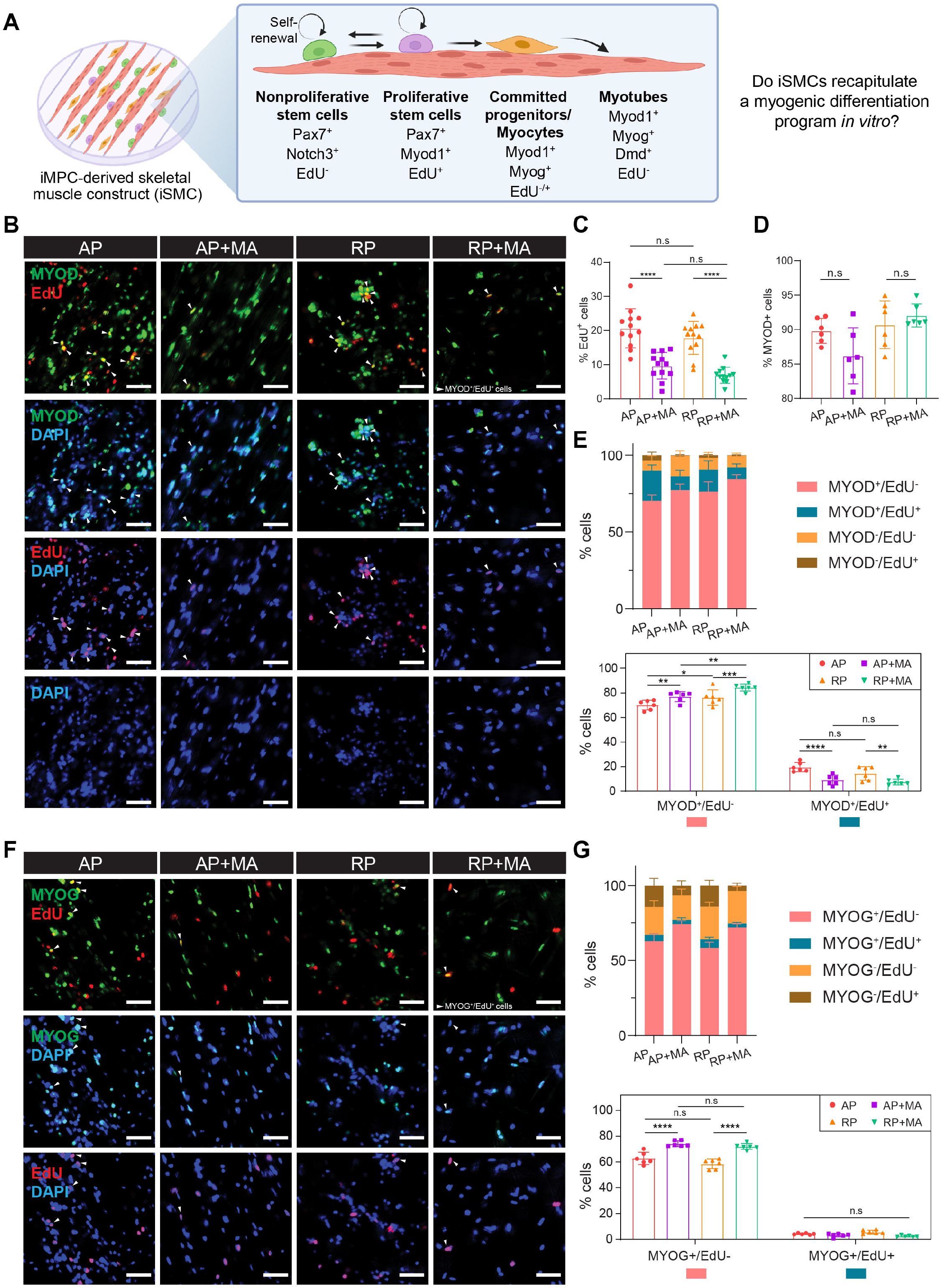
Proliferation and myogenic differentiation analysis of iSMCs. **(A)** An illustration of putative cell populations present in bioengineered iSMCs. **(B)** Representative immunofluorescence images of iSMCs stained for MYOD (green), EdU (red), and DAPI (blue) for the indicated conditions. Scale bar, 50 µm. **(C)** Graph showing quantification of EdU positive cells. The data are shown as means□±□S.D (n=12, 3 different images from 4 independent experiments were quantified). Statistical significance was determined by one-way ANOVA (****p<0.0001, n.s = nonsignificant). **(D)** Quantification of MYOD positive cells shown in (B). The data are shown as means□±□S.D (n=6, 3 different images from 2 independent experiments were quantified). Statistical significance was determined by one-way ANOVA (n.s = nonsignificant). **(E)** Quantification of cells in the indicated condition shown in (B). The data are shown as means□±□S.D (n=6, 3 different images from 2 independent experiments were quantified). Statistical significance was determined by two-way ANOVA (*p<0.5, **p<0.01, ***p<0.001, ****p<0.0001, n.s = nonsignificant). **(F)** Representative immunofluorescence images of iSMCs stained for MYOG (green), EdU (red), and DAPI (blue) for the indicated conditions. Scale bar, 50 µm. **(G)** Graph showing quantification of cells corresponding to the indicated condition shown in (F). The data are shown as means□±□S.D (n=6, 3 different images from 2 independent experiments were quantified). Statistical significance was determined by two-way ANOVA (*p<0.5, ****p<0.0001, n.s = nonsignificant).

To further corroborate this analysis, we performed immunostaining for the differentiation marker MYOG in concert with an EdU analysis (Figure 3F). In agreement with the immunostaining for MYOD, iSMCs in all conditions had approximately 60% MYOG^+^/EdU^-^ differentiated cells, while MA incorporation increased the number of non-proliferating MYOG^+^ cells compared to their respective conditions without MA (Figures 3G and S2B). Of note, a small percentage of MYOG^+^/EdU^+^ cells were detected across all conditions, in accordance with a previous study that reported proliferation of MYOG-expressing mononucleated cells (Figures 3G and S2C)^44^. In summary, the quantitative analyses revealed several myogenic cell populations in iSMCs under all conditions, without major differences between aligned or randomly patterned iSMCs. However, fewer proliferating cells were detected in the presence of MA, suggesting enhanced terminal differentiation when using this biomaterial. This observation is in agreement with the higher fusion index documented in MA-treated iSMCs, as well as reduced number of PAX7^+^ stem cells.

### Deconstructing cellular heterogeneity of iSMCs by scRNA-seq

Our analyses thus far revealed distinct cell populations in iSMCs consisting of stem, progenitor and differentiated cells. To further characterize the transcriptional heterogeneity of aligned iSMCs as well as the potential effect of MA supplementation in depth, we conducted scRNA-seq analysis of iSMCs under AP or AP+MA conditions (Figure 4A). First, the cells were collected from iSMCs by enzymatic digestion, and then subjected to 10x platform processing and sequencing after removal of multinucleated myotubes by filtration (Figure 4A). The sequencing data were processed individually using the Seurat v4.2.1 pipeline^45^ for each condition and integrated together as represented by a unified UMAP (Figure 4B). Notably, 5 cell clusters (C) were detected in the integrated UMAP and no differences were observed in the cell type composition between AP and AP+MA conditions (Figures 4B, 4C and S3A). These 5 cell clusters were annotated as two stem cell populations that were positive for *Pax7* (C1, C2), a committed progenitor cell population expressing *Dll1* (C3) and two differentiated cell populations expressing *Myog* (C4, C5) (Figure 4B).

**Figure 4.**
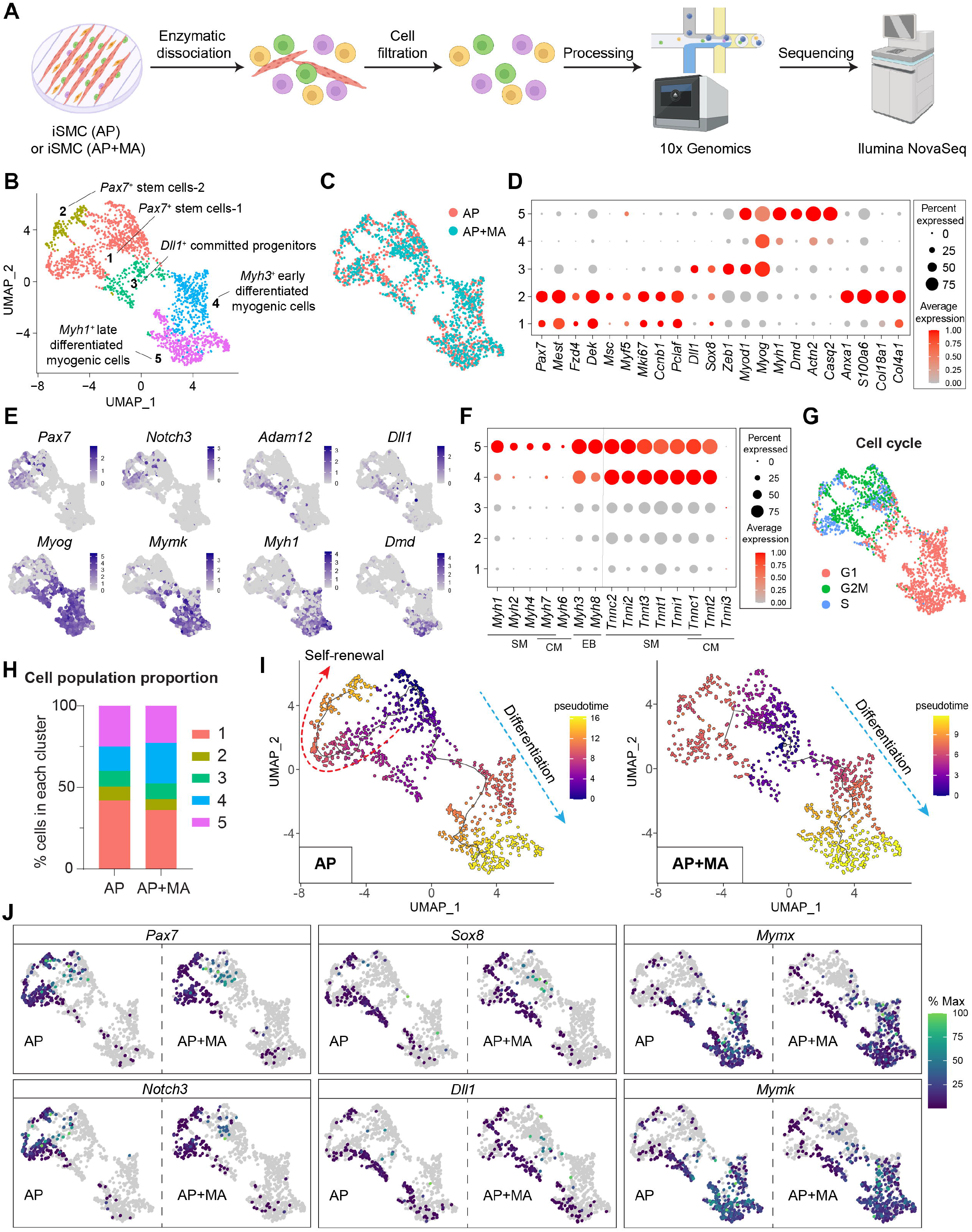
scRNA-seq analysis of iSMCs. **(A)** Graphical schematic of the experimental design to analyze the cell populations present in iSCMs. **(B)** UMAP projection showing the integrated objects of AP and AP+MA conditions, denoting 967 and 989 cells extracted for scRNA-seq analysis of iSMCs under the AP or AP+MA condition, respectively. Cells are colored by clusters. **(C)** UMAP projection of integrated scRNA data sets colored by condition. **(D)** Dot plot showing the expression level of the indicated marker genes for each cell cluster. **(E)** Feature plots showing the expression level of the indicated genes. **(F)** Dot plot showing the expression level of the indicated genes that are associated with skeletal muscle (SM), cardiac muscle (CM) and embryonic muscle (EB) markers. **(G)** UMAP projection of integrated scRNA data sets colored by each cell cycle state. **(H)** Bar graph showing the cell cluster distribution in percentages for the two examined iSMCs. **(I)** Single cell pseudotime trajectory analysis of the two examined iSMCs. Cells are colored by pseudotime values. **(J)** UMAP showing the expression of the indicated genes along the trajectory shown in (I).

Next, a closer investigation into the two stem cell clusters C1 and C2 revealed similar expression of satellite cell-associated genes including *Pax7, Msc, Myf5* and *Heyl*: however, C2 further exhibited expression of extracellular matrix-related genes including *Col18a1* and *S100a6*, as previously observed for a *Pax7*^+^ subpopulation in an established iMPC clone (Figures 4D, 4E S3B and S3C)^41^. The committed progenitor cell population (C3) expressed *Dll1, Sox8* and *Zeb1* (Figures 4D, 4E and S3B). In our former study, we reported on a transitory cell cluster expressing *Dll1* that interacted with *Notch*^+^ muscle stem cells via ligand-receptor binding^41^. Its detection in iSMCs supports our prior assumption that this cell population represents a nexus between myogenic stem and differentiated cells in iMPCs. In respect to the differentiated cell clusters C4 and C5, both expressed elevated levels of *troponin* isoforms and embryonic *Myh* isoforms such as *Myh3* and *Myh8*, yet C5 exhibited a higher expression of terminally differentiated markers including fast- and slow-twitching *Myh* isoforms (*Myh1, Myh4, Myh*7), dystrophin (*Dmd), Actn2* and *Casq2* (Figure 4F). This observation implies that C5 is at the tail end of the myogenic differentiation program, denoting cells that have exited the cell cycle prior to cell fusion. In support of this finding, a cell cycle analysis revealed a high expression of cell cycle regulators indicative of a proliferation state (S and G2M) in the stem and progenitor cell populations C1-3, and a high expression of non-proliferating cell cycle regulators (G1) in the differentiated cell populations C4-5 (Figure 4G). Although we did not detect prominent differences between the conditions AP and AP+MA, the cell fraction of C1 was higher under the AP condition (41.9% in AP vs. 36.0% in AP+MA), whereas that of C4 was higher under the AP+MA condition (14.8% in AP vs. 25.0% in AP+MA) (Figure 4H). As C1 is enriched in stem cell markers such as *Pax7, Heyl* and *Fzd4*, whereas C4 is enriched in differentiation markers such as *Myog*, this analysis supports our prior observation that MA supplementation reduces the number of stem cells while augmenting differentiation capacities.

To gain further insights into the gene expression dynamics during myogenic differentiation in iSMCs, we performed a pseudotime trajectory analysis. This effort revealed that the initiation of the myogenic differentiation program under the AP condition starts with C1 and bifurcates into either C2 or C3-5 arms, whereas under AP+MA the initiation similarly starts at C1 however predominantly proceeds to the C3-5 arm (Figure 4I). Accordingly, *Pax7* and *Notch3* expression during the transition from C1 to C2 were predominantly upregulated in the AP condition, whereas a similar kinetic progression was observed for the committed progenitor markers (*Sox8* and *Dll1*) and differentiated markers (*Mymx* and *Mymk*) under both conditions (Figure 4J). This observation cautiously implies an impaired self-renewal in the stem cell populations C1 and C2 under the AP+MA condition. However, a confirmation of this analysis would require further investigation to rule out the possibility that this effect might be derived from technical limitations due to low gene/UMI counts under the AP+MA condition. In summary, using scRNA-seq analysis we unveiled the cell populations comprising iSMCs, dissecting the different stages of the myogenic differentiation program. Overall, no major transcriptional differences were detected between the two examined conditions. However, subtle differences between AP and AP+MA conditions were documented for certain cell populations, pointing towards an increase in self-renewal of PAX7^+^ stem cells under the AP condition, and an augmented myogenic differentiation propensity under the AP+MA condition.

### Assessing the regeneration potential of iSMCs in an injury-based model

Our results thus far revealed that iSMCs contain proliferating PAX7^+^ stem cells that can robustly differentiate into aligned myotubes. This raises the interesting question whether the iMPC-derived biomimetic constructs can regrow aligned myotubes following an ectopic insult, similar to muscle regeneration *in vivo*. To investigate this hypothesis, we treated iSMCs with cardiotoxin (CTX), a widely used myonecrotic agent that can trigger extensive damage to muscle fibers or myotubes but rarely to satellite cells^46,47^ (Figure 5A). We chose to analyze iSMCs across various time points upon CTX administration, which occurred on the day of cell seeding, as well as after CTX withdrawal (Figure 5A). At 24 hrs post CTX administration, a few MYHC^+^ myotubes were detected under either the AP or AP+MA condition, although many PAX7^+^ cells were observed (Figure 5B). Accordingly, we noted that CTX administration increased the expression level of *Pax7* and the proliferation marker *Mki67* whereas it decreased the expression level of differentiation related genes such as *Myf6* and *Myh1* under both conditions, most likely due to the depletion of differentiated skeletal muscle cells (Figure 5C). Two days post CTX administration, we analyzed the constructs and observed that multinucleated MYHC^+^ myotubes were predominantly formed under the AP condition, growing in thickness at days 4 and 8 (Figure 5D). Accordingly, more PAX7^+^ cells were observed under the AP condition at the analyzed time points (Figure 5D). At day 8 post CTX administration, large multinucleated myotubes were detected under the AP condition, together increased presence of PAX7^+^ stem cells (Figure 5D).

**Figure 5.**
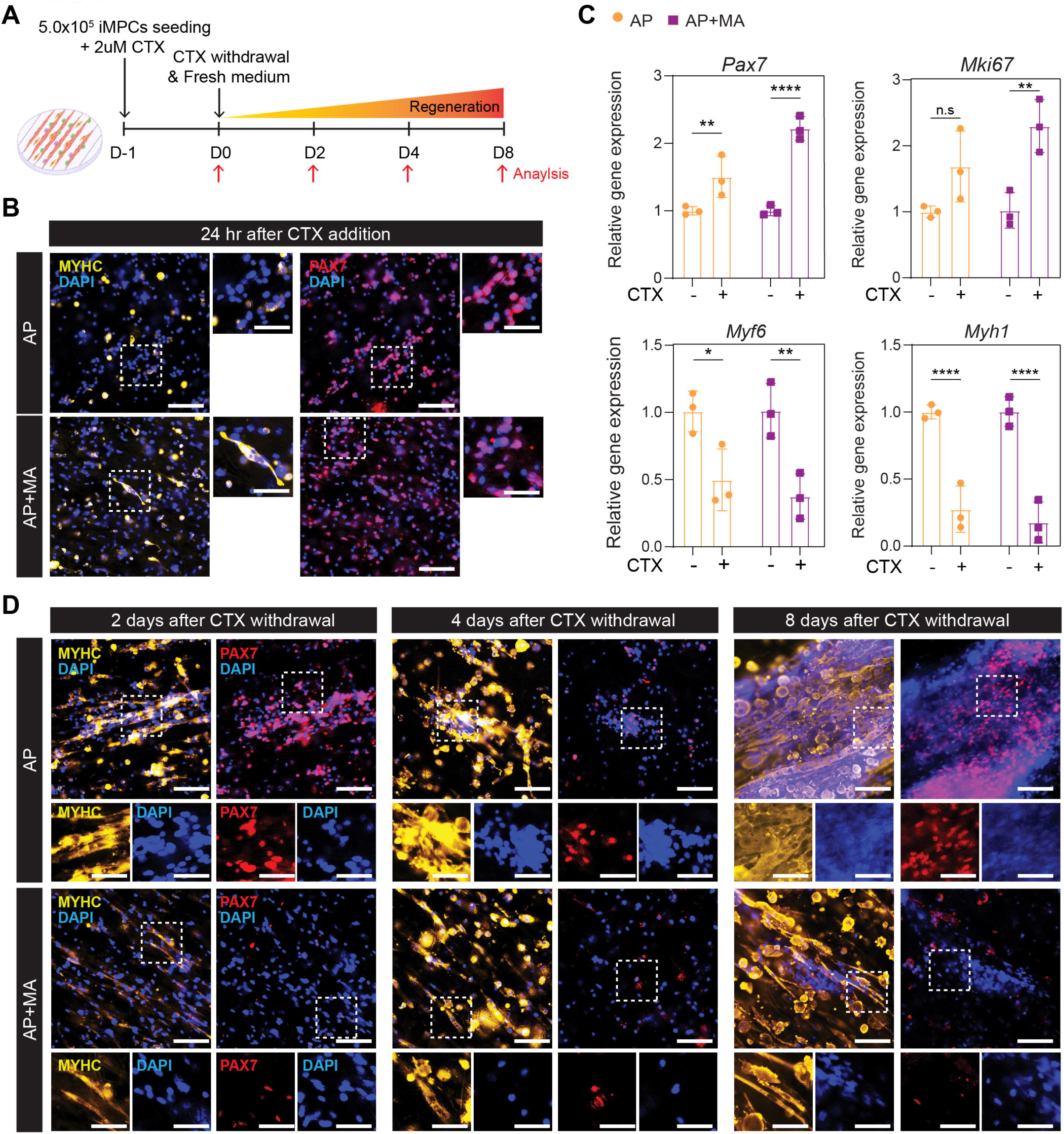
Analysis of iSMCs subjected to a myonecrotic agent. **(A)** Graphical schematic illustrating the indicated time points used to evaluate the regeneration capacity of iSMCs following a cardiotoxin (CTX)-induced myofiber damage. **(B)** Representative immunofluorescence images of iSMCs stained for MYHC (yellow), PAX7 (red), and DAPI (blue) in the respective conditions. The images were taken 1 day after CTX treatment. Scale bar, 100 µm. Scale bar of zoom-in images, 50 µm. **(C)** qRT-PCR analysis of the indicated genes that are associated with a myogenic stem or differentiated cell fate, as well as the proliferation marker *Mki67*. The data are shown as means□±□S.D (n=3 independent experiments). Statistical significance was determined by a one-way ANOVA (*p<0.5, **p<0.01, ****p<0.0001). **(D)** Representative immunofluorescence images of iSMCs stained for MYHC (yellow), PAX7 (red), and DAPI (blue) in the indicated conditions. Scale bar, 100 µm. Scale bar of zoom-in images, 50 µm.

Given the abundant PAX7^+^ stem cells observed after CTX exposure in the AP condition, we subjected iSMCs to a 2^nd^ CTX administration to assess their regeneration competency in a serial injury model. Under this experimental design, the 2^nd^ CTX treatment was higher in concentration and applied to the iSMCs 4 days after recovery from the 1^st^ CTX treatment, and the respective analysis was performed up to 5 days after the second injury (Figure 6A). Similar to the 1^st^ CTX treatment, another round of injury resulted in extensive damage and destruction of myotubes under both conditions (Figure 6B). Importantly, the AP+MA condition poorly regenerated myotubes after this treatment, whereas the AP condition successfully regenerated aligned iSMCs consisting of PAX7^+^ stem cells and MYHC^+^ myotubes 5 days after reinjury (Figure 6C). This observation suggests that self-renewing PAX7^+^ stem cells are important for regenerating myotubes upon CTX-induced muscle damage, and further corroborated the scRNA-seq analysis which demonstrated that a cycling PAX7^+^ stem cell population gives rise to differentiated muscle cells in iSMCs. In summary, following a single or a serial CTX injury, self-renewing muscle stem cells were able to regenerate aligned myotubes in iSMCs in the absence of MA, resembling myofiber regeneration *in vivo*.

**Figure 6.**
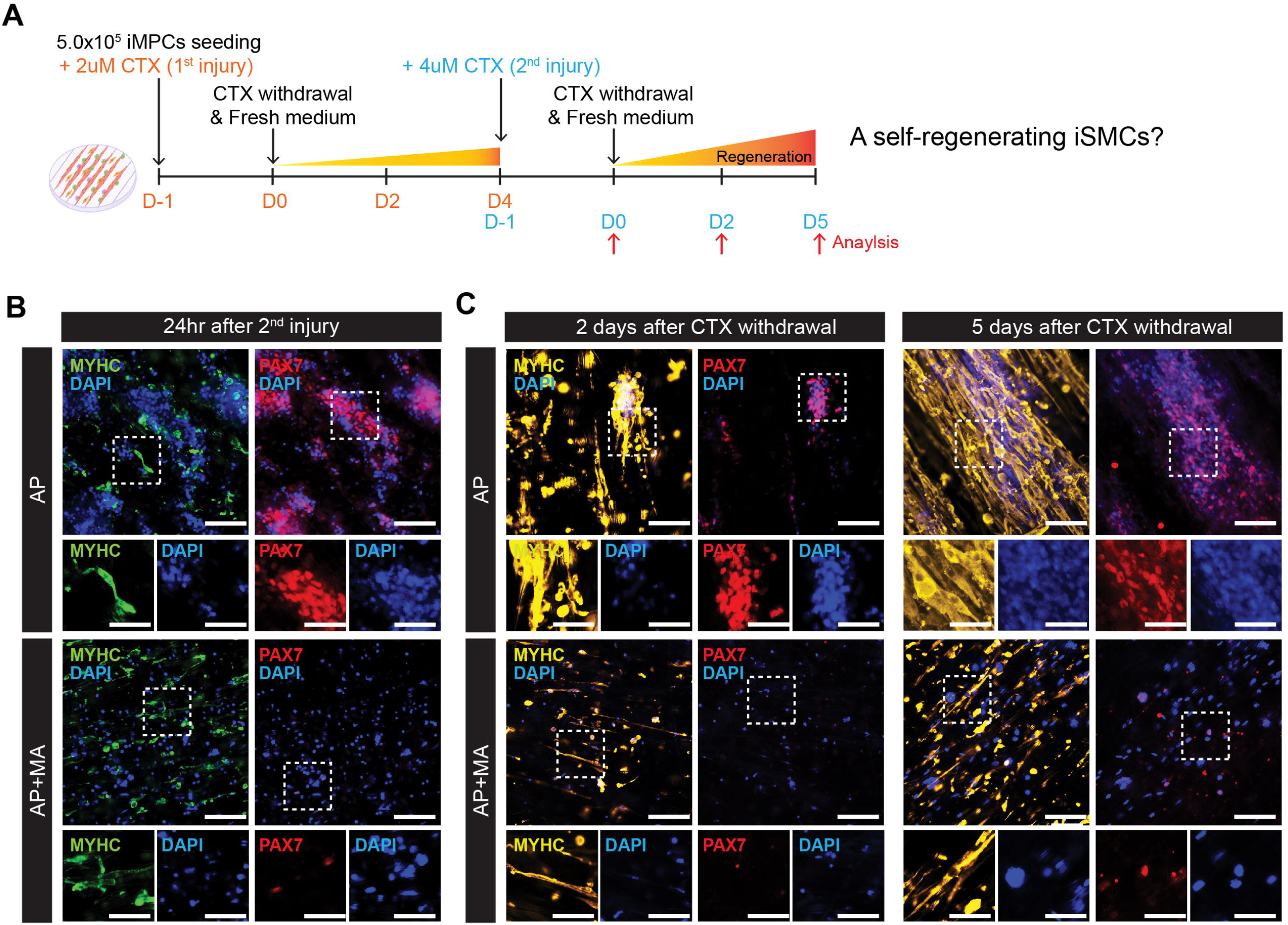
Analysis of iSMCs subjected to a serial injury assay. **(A)** Graphical schematic illustrating the experimental procedure used to assess the regeneration capacity of iSMCs after reinjury with CTX. **(B)** Representative immunofluorescence images of iSMCs stained for MYHC (green), PAX7 (red), and DAPI (blue) in the respective indicated conditions. The images were taken 1 day after the 2^nd^ CTX treatment. Scale bar, 100 µm. Scale bar of zoom-in images, 50 µm. **(C)** Representative immunofluorescence images of iSMCs undergoing regeneration after a 2^nd^ CTX injury and stained for MYHC (yellow), PAX7 (red), and DAPI (blue) for the indicated conditions. The images were taken 2 or 5 days after CTX withdrawal. Scale bar, 100 µm. Scale bar of zoom-in images, 50 µm.

## Discussion

In this study, we established a biomimetic skeletal muscle construct by seeding iMPCs on electrospun PCL substrates, rendering myotube alignment in conjunction with self-renewing muscle stem cells. Furthermore, incorporation of MA enhanced the differentiation capacity of iMPCs in the scaffolds, while mitigating their self-renewal. We further dissected the cell populations in iSMCs using a variety of molecular assays, and demonstrated their capacity to regenerate aligned myotubes in a single or serial injury model.

Several implications extend from our study. In respect to the use of MA, we envision it entails both advantageous and disadvantageous aspects for tissue engineering purposes. The MA is an intricate and undefined protein and macromolecule matrix that is widely used for conventional myoblast culture, as it enables satellite cell attachment to plastic and supports proliferation in medium containing high serum^8,48^. However, for the system reported in this study, MA induced morphological and cellular changes in iSMCs, manifesting increased differentiation and mitigating self-renewal of muscle stem cells, most likely due to signaling molecules present in MA such as IGF1 and TGF-β^49,50^. As such, MA supplementation might be useful for depleting PAX7^+^ stem cells and enhancing myotube formation in the event predominantly differentiated cells are desired in iSMCs. Furthermore, we observed by scRNA-seq analysis that MA supplementation was subtle, affecting primarily two cell populations. It is important to note however that scRNA-seq captures only the mononucleated fraction of iSMCs, and as such conducting single nucleus RNA-seq analysis of multinucleated myotubes might lend additional insights into the effect of MA supplementation on the maturation state of iSCMs.

To date, a multitude of studies reported aligned myotube formation in 3D for tissue engineering purposes^9-11,33^. In respect to the method used in this study, the advantages of using PCL is its biocompatibility, easy modification and FDA-approval^51^. Additionally, PCL does not include cell adhesion proteins that might affect the interactions between the cell populations comprising iMPCs, which are highly heterogeneous and require expansion in the presence of a small molecule cocktail that modulates several signaling pathways^37^. Given that PCL is also biodegradable, the use of iSMCs is attractive for *in vivo* therapeutic approaches aiming to treat extensive muscle degeneration in the form of late-stage Duchenne muscular dystrophy (DMD) or volumetric muscle loss (VML)^52^.

Another advantage of the approach reported herein is the unique use of iMPCs as a cell source to engineer skeletal muscle. As of today, only one study reported using directly reprogrammed myogenic progenitors for tissue engineering purposes, showcasing their potential to treat VML^52^. Similarly, we believe the use of iMPCs was key to rapidly producing iSMCs, as solely iMPC seeding in presence of high serum and F/R/C supplementation for as little as 2 days was required. This is a consequence of the intrinsic attributes of iMPCs, including rapid muscle stem cell proliferation in conjunction with robust differentiation into myotubes, unlike most conventional myoblast cell lines that are cultured either under proliferation or differentiation conditions^41^. For example, the most widely used immortalized myogenic cell line C2C12 is typically employed only for its fusion attributes^53^. However, C2C12 are karyotypically abnormal, immortalized and lack primary satellite-like muscle stem cells, representing a major limitation^7,53,54^.

In comparison to iMPCs cultured on plastic, iSMCs represent a more organized model, yet they do not fully mirror an *in vivo* skeletal muscle tissue which contains a variety of cell types including QSCs that are dormant under homeostatic conditions^55^. In contrast, iSMCs appear highly proliferative, mirroring an ongoing regeneration response *in vitro*, most likely due to the presence of the F/R/C small molecule cocktail. It will be of interest to assess if a fraction of the Pax7^+^ cells in iSMCs exit the cell cycle and return to quiescence, as previously reported for SCs seeded on myofibers^56-58^. Additionally, it will be of interest to assess whether serum withdrawal or modulation of additional signaling pathways might confer quiescence rather than activation on the Pax7^+^ cell population in iSMCs, rendering them more similar to muscle tissue during homeostasis^57,59,60^.

Constructing an intact skeletal muscle tissue *in vitro* is a major challenge, as in addition to satellite cells and muscle fibers skeletal muscle tissue contains a plethora of additional resident cells including endothelial cells, fibro-adipogenic progenitors, immune cells and motor neurons^55^. It will be of interest to seed these cell types into iSMCs to generate a more physiologically relevant construct that might better mimic cellular attributes and interactions present in muscle tissue *in vivo*. Furthermore, it will be of interest to assess whether contraction in iSMCs can be controlled by optogenetic, chemical or electrical stimuli as previously shown for other myogenic cell types ^34,47,61^. Finally, an attractive utility for iSMCs is to model muscular dystrophies such as DMD *in vitro*, potentially replacing the use of animal models and supporting the “Replace, Reduce, Refine” (3R) animal experimentation guidelines. To date, several studies reported on fabricating 3D engineered skeletal muscle constructs using DMD patient-derived myoblasts^62^ or iPSC-derived myogenic precursor cells^30,63^. Recently, our lab developed an *in vitro* cellular model for DMD by reprogramming fibroblasts into iMPCs that do not express dystrophin, enabling to restore dystrophin expression *in vivo* after genetic correction with CRISPR-Cas9^64^. We envision that DMD iSMCs could be employed to investigate disease pathology *in vitro*, or as a platform to test therapeutic interventions that might be applicable to human DMD patients.

## Acknowledgments

We wish to thank Xhem Qabrati, Giada Bacchin and Ajda Lenardic for their constructive comments and feedback. We are further grateful to the Functional Genomics Center Zurich (FGCZ) facility for their help with sample processing and sequencing.

## Funding

This work was supported by an Eccellenza Grant of the Swiss National Science Foundation, PCEGP3_187009 (OBN), as well as a European Union’s Horizon 2020 research and innovation program under the Marie Skłodowska-Curie grant agreement, 812765 (SJF).

### Author contributions

Conceptualization: IK, SL, OBN, SJF

Formal analysis: IK, SL, AG

Funding Acquisition: OBN, SJF

Investigation: IK, SL

Methodology: IK, SL, AG

Supervision: OBN, SJF

Visualization: IK, SL

Writing – original draft: IK, SL, OBN

Writing – review & editing: IK, SL, AG, SJF, OBN

## Data and materials availability

The scRNA-Seq datasets reported in this study are available at Gene Expression Omnibus (GEO) repository (GSE222162).

## Material and Methods

### Fabrication of PCL scaffolds using electrospinning

To produce electrospun PCL scaffolds, we used PCL (Sigma-Aldrich, 440744, Mn=80,000 g mol^-1^) that was dissolved for 24 hrs under constant stirring at 10% (w/v) in a 4:1 chloroform: ethanol mixture. Electrospinning was performed using a commercially available electrospinning machine (IME Technologies, EC-CLI) for 60 mins using the following parameters and conditions: electric potential of the nozzle = 15 kV, electric potential of the collector = -2 kV, size of needle= 22G, flow rate = 7.5 mL/h, distance between nozzle and collector = 15 cm and mandrel rotation speed = 100 RPM or 1500 RPM depending on the scaffold group. Electrospun nanofibers were collected onto a ø 90×180 mm cylindrical mandrel at room temperature. After spinning, the membranes were cut into cylinders with a ø 8 mm biopsy punch.

### Surface modification and sterilization of PCL scaffolds

PCL membranes were treated with 1M NaOH for 4 hrs, followed by several washes with PBS and sterilization in 70% Ethanol overnight. The scaffolds were then washed with PBS in sterile conditions, placed under UV lamp for 20 mins and used for cell seeding.

### Fiber alignment angle and fiber size measurement

As a first step, the fibrous structure images comprising the electrospun scaffolds were captured by FE-SEM. Next, acquired images were analyzed using the ImageJ software (NIH, United States) to measure the alignment angle and fiber diameter. The alignment angle of electrospun fibers was measured by using the “Angle Tool” function in ImageJ which measures the angles between the pattern direction and the longest axis of the fiber. For the fiber diameter measurement, the “Set Scale” function in ImageJ was used first to set unit length by converting the obtained length from the scale bar in FE-SEM image. Then, “Straight line” function in ImageJ was used to measure the fiber diameter from the image.

### Cell media composition

The iMPC medium contained KnockOut (KO) DMEM (Thermo Fisher Scientific, 10829018) supplemented with 10% KO Serum Replacement (Thermo Fisher Scientific, 10828028), 10% FBS (Thermo Fisher Scientific, 10270106), 1% GlutaMAX (Thermo Fisher Scientific, 35050061), 1% non-essential amino acids (Thermo Fisher Scientific, 11140050), 1% Pen-Strep (Thermo Fisher Scientific, 15140122), 0.1% 2-Mercaptoethanol (Thermo Fisher Scientific, 21985023), 10 ng/mL basic FGF (R&D Systems, 233-FB), 5 µM Forskolin (F) (R&D Systems, 1099/50), 5 µM RepSox (R) (R&D Systems, 3742/50) and 3 µM CHIR99021 (C) (R&D Systems, 4423/50).

### Generation of iSMCs

Reprogramming of fibroblasts into the iMPCs used in this study has been previously described^41^. For iSMC production, about 5×10^5^ or 1.0×10^5^ iMPCs were trypsinized and resuspended in 10µl iMPC medium or 4% Matrigel (Corning, 356237) diluted in low glucose DMEM (Thermo Fisher Scientific, 31885023). Next, resuspended iMPCs were seeded on top of the sterile PCL scaffolds and placed in a cell culture incubator (37°C, 5% CO2) for 3-4 hrs until cells were attached to the scaffolds. The iMPC medium was then added to each well and changed freshly every other day. All subsequent analyses (excluding the CTX injury model) were conducted 2 days after cell seeding.

### Field Emission Scanning Electron Microscopy (FE-SEM)

For structure observation, scaffolds were cut into ø 8 mm using a biopsy punch and fixed on metal stubs with a carbon tape coated with platinum/palladium (80/20) sputtering (Safematic, CCU-010). Field Emission-Scanning Electron Microscopy (FE-SEM) (Hitachi, SEM SU5000) was used to capture the surface topology, fiber alignment and microstructure of the scaffolds at 5kV. For cell attachment, 5.0×10^4^ iMPCs were seeded onto the scaffolds and cultured for 2 days in iMPC medium. The cells were then washed with PBS and fixed with 4% paraformaldehyde (PFA) and 3% glutaraldehyde (both suspended in PBS). As the next step, the cells were fixed with 2% osmium tetroxide in PBS (Polysciences, 23310-10) for membrane preservation, and dehydrated with a graded ethanol series ending in absolute ethanol. After dehydration, the cells were washed with hexamethyldisilazane (Sigma-Aldrich, 440191). Finally, the samples were mounted and coated with platinum/palladium (80/20) sputtering for FE-SEM.

### Immunofluorescence staining

At defined time points, iSMCs were fixed with 4% paraformaldehyde (Alfa Aesar, 43368) for 10 mins and washed with PBS twice. To avoid random binding of antibodies and to permeabilize the cell membrane, iSMCs were incubated for 30 mins in blocking solution containing 2% BSA (AppliChem, 9048-46-8) and 0.5% Triton™ X-100 (Sigma-Aldrich, 9002-93-1). The blocking solution was then discarded and iSMCs were incubated for 2 hrs with primary antibodies diluted in blocking solution, followed by washing and addition of secondary antibodies and 4′,6-diamidino-2-phenylindole (DAPI) (1:1000, Thermo Fisher Scientific, 62248) for 30 mins. The primary antibodies used in this study include anti-Mouse Myosin Heavy Chain (1:1000, R&D Systems, MAB4470), anti-Human/Mouse/Rat/Chicken Pax7 (5μg/ml, R&D Systems, MAB1675), anti-Human/Mouse MyoD (5.8A) (1:200, Thermo Fisher Scientific, MA512902) and anti-myogenin (F5D) (1:250, Santa Cruz, sc-12732). The secondary antibodies used in this study were Goat Anti-Mouse IgG2b (Alexa Fluor 546) (1:500, Thermo Fisher Scientific, A21143), Goat Anti-Mouse IgG1 (Alexa Fluor 647) (1:500, Thermo Fisher Scientific, A21240) and Goat Anti-Mouse IgG1 (Alexa Fluor 546) (1:500, Thermo Fisher Scientific, A21123). Fluorescence images were taken using either a confocal (Carl Zeiss, LSM 880 airyscan) or an inverted microscope (Nikon ECLIPSE Ti2).

### Quantitative real-time PCR (RT-qPCR)

The iSMCs were placed in 1.5ml Eppendorf tubes and directly subjected to RLT cell lysis buffer supplied in the RNeasy kit (Qiagen, 74104) for 1 min while vortexing. According to the manufacturer’s instruction, the cell lysates were then used for RNA extraction and converted into cDNA by a High-Capacity cDNA Reverse Transcription Kit (Thermo Fisher Scientific, 4368814). The same amount of cDNA for each sample was then used for probe-based RT-qPCR. The probes used in this study were purchased from Integrated DNA Technologies (IDT, Coralville, USA) and include murine(m)*Gapdh* (Mm.PT.39a.1), m*Pax7* (Mm.PT.58.12398641), m*Notch3* (Mm.PT.58.7794053), m*Heyl* (Mm.PT.58.17090120), m*Myod1* (Mm.PT.58.8193525), m*Myog* (Mm.PT.58.30712483.gs), m*Myf6* (Mm.PT.58.33344984) and m*Myh1* (Mm.PT.58.11712984). For *Mki67*, PowerUp SYBR Green Master Mix (Thermo Fisher Scientific, A25741) was used using the following primer sequences: forward primer 5’-CAGTTTGGCGACATTAGCAGA-3’ and reverse primer 5’-GCAACTATCTTGGCAACATCCTC-3’.

### EdU analysis

To detect proliferating cells in iSMCs, the Click-iT™ EdU Cell Proliferation Kit for Imaging, Alexa Fluor™ 647 dye (Thermo Fisher Scientific, C10340) was used according to the manufacturer’s protocol. Briefly, 10mM EdU was added to iMPC medium at a final concentration of 10µM and then applied to the iSMCs for 2 hrs at 37°C. Then, iSMCs were fixed with 4% PFA for 10 mins and incubated in blocking solution for 20 mins, followed by Click-iT™ reaction cocktail for 30 mins at RT for EdU color development and staining. To combine an EdU analysis with conventional immunostaining, primary and secondary antibodies were applied as described in the respective Immunofluorescence staining method section.

### scRNA-seq library construction

To construct scRNA-seq libraries we first detached the cells from the PCL scaffolds by treating them with 2mg/ml Dispase II (Thermo Fisher Scientific, 17105041) for 20 mins at 37°C, followed by 0.25% Trypsin-EDTA for 10 mins. Scaffolds were then removed, and remaining cells were centrifugated for 5 mins at 300g. To disaggregate iMPC clumps into single cells, 0.25% Trypsin-EDTA was again applied to the cell pellet for 5 mins and neutralized with iMPC medium. Cells were then filtered through 40 μm strainer (VWR, 734-0002) and resuspended in iMPC medium and immediately used for scRNA-seq library production. The libraries were constructed using Single cell 3’ reagent kit v3.1 on 10x platform (10x Genomics, Pleasanton, CA) as detailed in a previous study^41^. Libraries were sequenced in full SP flow cell of NovaSeq with paired-end 28-91bp.

### Analysis of scRNA-seq data

scRNA-seq data were demultiplexed and mapped against the mouse reference genome assembly GRCm39 (GENCODE release M26) using 10x Genomics CellRanger v7.0.0^65^. Downstream analysis of the resulting filtered feature-barcode count matrices was performed using the Seurat v4.2.1 pipeline^45,66^. For quality control, cells with unique feature counts < 250 and > 9,000, and mitochondrial gene counts > 10% were removed. For each condition, filtered data were log normalized, scaled and the top 2,000 variable features were detected. Samples were integrated based on feature anchors^45^. Integrated data were scaled, and principal component analysis (PCA) was performed using the highly variable features for dimensional reduction. Cells were clustered based on first 30 principal components (PCs) using Louvain algorithm^67^ with a resolution of 0.5. Clustered cells were visualized in two-dimensional space using the uniform manifold approximation and projection (UMAP)^68^ of same PCs. For each cell cluster, conserved marker genes were identified based on the Wilcoxon rank-sum test with log2 fold-change > 0.25 and adjusted p-value < 0.05 cut-offs. Unsupervised single cell trajectory analysis was conducted using Monocle3 v1.2.9^69,70^ and cells were ordered in pseudotime along the learned trajectory at the conditional level with cluster 1 functioning as the root.

### Cardiotoxin injury

To induce muscle injury in iSMCs, 5.0×10^4^ iMPCs were seeded onto PCL scaffolds and incubated in iMPC medium containing 2 μM cardiotoxin (CTX) (Latoxan, L8102) 4 hrs after cell attachment. On the next day, iMPC medium containing CTX was discarded, and fresh medium was applied. For the 2^nd^ injury, 4 days after the 1^st^ CTX removal, iSMCs were treated with 4μM CTX in iMPC medium for 24 hrs and then incubated in fresh medium for recovery.

### Statistical analysis

We conducted statistical analysis using the software GraphPad Prism as indicated in each figure legend.

