## Supplementary information for "A self-renewing biomimetic skeletal muscle construct engineered using induced myogenic progenitor cells"

### Figure S1

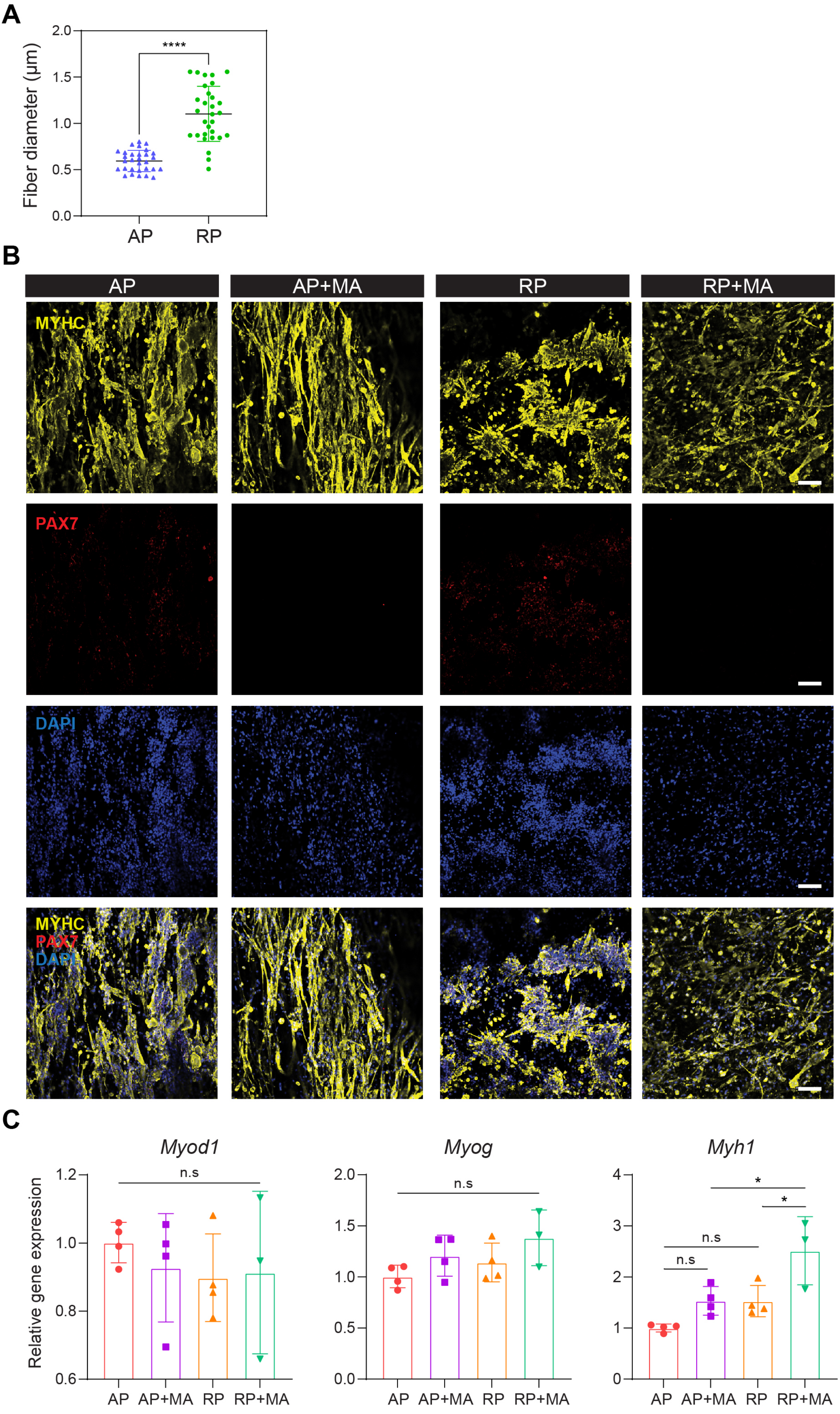

A
Fiber diameter (μm)
2.0
1.5
1.0
0.5
0.0
****
AP
RP
B
AP
AP+MA
RP
RP+MA
MYHC
PAX7
DAPI
MYHC
PAX7
DAPI
C
Myod1
Myog
Myh1
Relative gene expression
1.2
1.0
0.8
0.6
2.0
1.5
1.0
0.5
0.0
4
3
2
1
0
n.s
n.s
n.s
n.s
*
*
AP
AP+MA
RP
RP+MA
AP
AP+MA
RP
RP+MA
AP
AP+MA
RP
RP+MA

**Figure S1.**
**(A)** Diameter measurement of electrospun PCL nanofibers. The data are shown as means ± S.D (n=30). Statistical significance was determined by a two-tailed unpaired t-test (****p<0.0001). **(B)** Representative immunofluorescence images of iSMCs stained for MYHC (yellow), PAX7 (red), and DAPI (blue) in the indicated conditions. Scale bar, 100 µm. **(C)** Graphs denoting qRT-PCR analysis of the indicated myogenic genes. The data are shown as means ± S.D (n=3-4 independent experiments). Statistical significance was determined by one-way ANOVA (*p<0.5, n.s = nonsignificant).

Figure S2

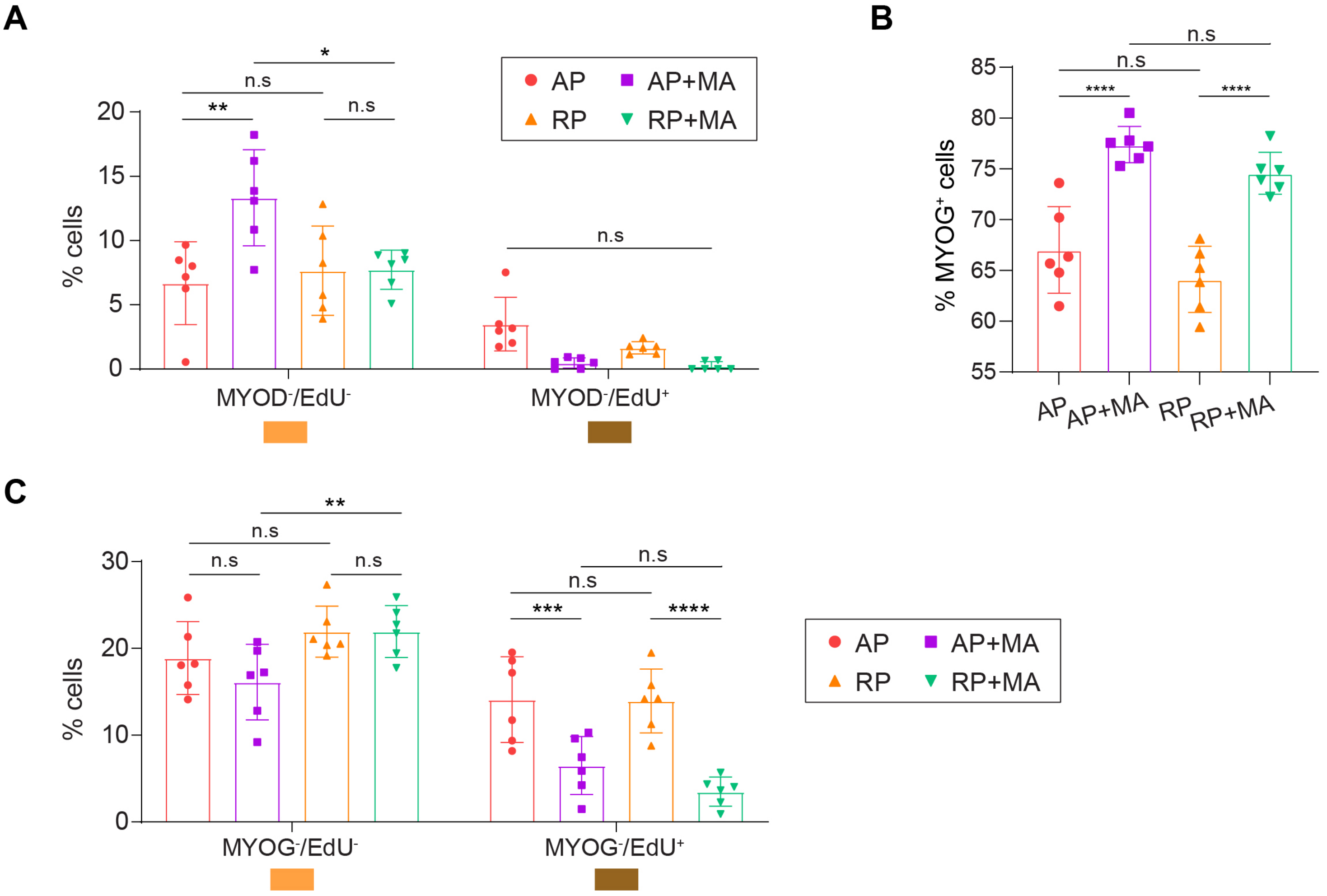

A
AP
AP+MA
RP
RP+MA
% cells
20
15
10
5
0
**
n.s
*
n.s
n.s
MYOD-/EdU-
MYOD-/EdU+
B
% MYOG+ cells
85
80
75
70
65
60
55
n.s
n.s
****
****
AP
AP+MA
RP
RP+MA
C
% cells
30
20
10
0
**
n.s
n.s
n.s
n.s
n.s
***
****
MYOG-/EdU-
MYOG-/EdU+
AP
AP+MA
RP
RP+MA

**Figure S2.**

**(A)** Graphs showing quantification of the images presented in Figure 3B. The data are shown as means ± S.D (n=6, 3 different images from 2 independent experiments were quantified). Statistical significance was determined by two-way ANOVA (*p<0.5, **p<0.01, n.s = nonsignificant). **(B)** Graph showing quantification of the MYOG positive cells shown in Figure 3F. The data are shown as means ± S.D (n=6, 3 different images from 2 independent experiments were quantified). Statistical significance was determined by two-way ANOVA (****p<0.0001, n.s = nonsignificant). **(C)** Quantification of the images shown in Figure 3F. The data are shown as means ± S.D (n=6, 3 different images from 2 independent experiments were quantified). Statistical significance was determined by two-way ANOVA (**p<0.01, ***p<0.001, ****p<0.0001, n.s = nonsignificant).

Figure S3

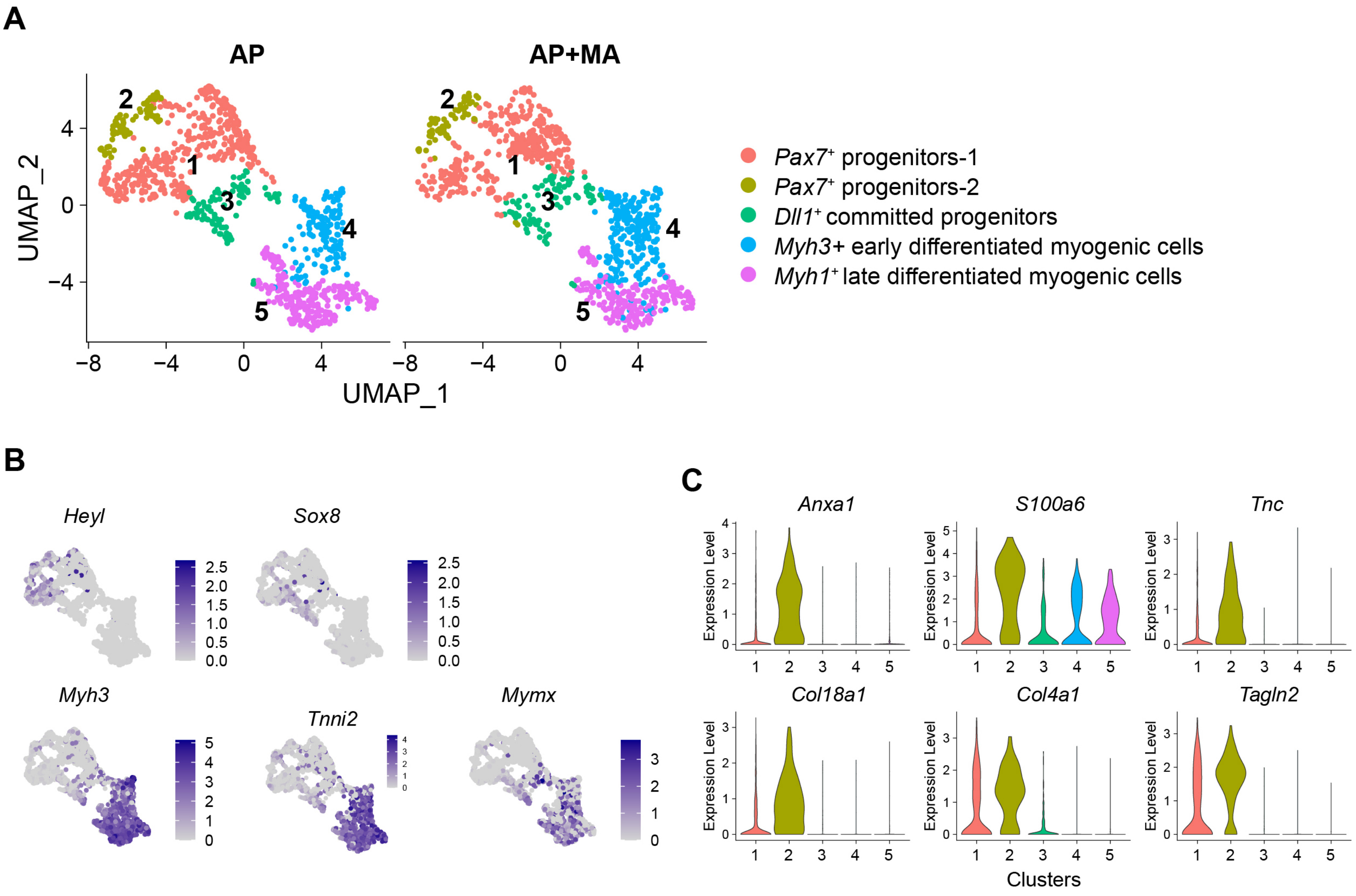

A
AP
AP+MA
UMAP_1
UMAP_2
Pax7+ progenitors-1
Pax7+ progenitors-2
Dll1+ committed progenitors
Myh3+ early differentiated myogenic cells
Myh1+ late differentiated myogenic cells
B
Heyl
Sox8
Myh3
Tnni2
Mymx
C
Anxa1
S100a6
Tnc
Col18a1
Col4a1
Tagln2
Expression Level
Clusters

**Figure S3.**

**(A)** UMAP projections based on scRNA-seq data of the individual conditions AP and AP+MA. Cells are colored by clusters. **(B)** Feature plots showing the expression level of the indicated genes. **(C)** Violin plots showing the expression level of extracellular matrix-related marker genes that are enriched in cluster 2.
